# The protein phosphatases MoPtc1 and MoPtc2 are induced during pathogen-host interactions and play synergistic roles in regulating MAPK pathways in *Magnaporthe oryzae*

**DOI:** 10.1101/2022.09.09.507255

**Authors:** Wilfred M. Anjago, Jules Biregeya, Mingyue Shi, Yixiao Chen, Yupeng Wang, Meilian Chen, Osakina Aron, Justice Norvienyeku, Wenyi Yu, Zonghua Wang, Zhang Dongmei

**Affiliations:** Ministry of Education Key Laboratory of Biopesticides and Chemical Biology, College of Life Science, Fujian Agriculture and Forestry University, Fuzhou, 350002, China; Institute of Oceanography, Minjiang University, Fuzhou, Minjiang University, Fuzhou, 350108, China

**Keywords:** *Magnaporthe oryzae*, Protein phosphatase, MAPK signaling pathway

## Abstract

Reversible protein phosphorylation is essential in cellular signal transduction. The rice blast fungus *Magnaporthe oryzae*, contains six putative type 2c protein phosphatases namely; MoPtc1, MoPtc2, MoPtc5, MoPtc6, MoPtc7 and MoPtc8 respectively. In this study, we carried out transcription expression analysis and found that MoPtc1, MoPtc2 and MoPtc7 are significantly induced during pathogen-host interactions. Subsequent deletions of MoPtc1 or MoPtc2 not only resulted in increased sensitivity to cell wall stress mediated by Congo red but also increased phosphorylation of Mps1-MAPK. By immunoblot analysis, we found that deletion of both MoPtc1 and MoPtc2 resulted in overstimulation of both the HOG1 and Pmk1 pathways in *M. oryzae*. We also demonstrate that MoPtc1 is recruited directly to Osm1 by the adaptor protein MoNbp2 to inactivate the Osm1 during hypoosmotic stress unlike in budding yeast. Finally, we show that type 2c protein phosphatases are localized in different cellular compartments in the life cycle of *Magnaporthe oryzae*. Taken together type 2C protein phosphatases MoPtc1 and MoPtc2 play synergistic roles in regulating MAPK signaling pathways in *M. oryzae*. This work expands our understanding of the MAPK signaling regulation circuits in *M. oryzae* and the essential roles of type 2C phosphatases in fine-tuning phosphorylation levels of MAPK during fungal development.

## Introduction

Most living organisms perceive and respond to environmental cues by signal transduction pathways involving Mitogen-activated Protein Kinases. In plants, these pathways mediate defence responses through the recognition of PAMPs (pathogen-associated molecular patterns) or MAMPs (microbe-associated molecular patterns) by pattern recognition receptors (PRRs), while in appressorium-forming pathogens, they trigger the development of infectious structures and promote infection by suppressing the plant defence system (1–4). The activation of MAPK signalling pathways occurs through phosphorylation-catalysed reactions orchestrated by protein kinases, while deactivation is achieved by the removal of a phosphate group from a serine, threonine or tyrosine residue of a MAPK. This reaction is coordinated by protein phosphatases in eukaryotes (5).

Protein phosphatases are broadly divided into two main groups, serine/threonine protein phosphatases and tyrosine protein phosphatases, based on their distinct structure, sequence homology and catalytic mechanism concerning substrate specificity (6, 7). Serine/threonine protein phosphatases are structurally distinguished into phosphoprotein phosphatases (PPPs), metal-dependent protein phosphatases (PPMs), and aspartate-based protein phosphatases (APPs) (8). Compared with other protein phosphatases, PPMs are monomeric enzymes whose catalytic subunit usually does not associate with its regulatory subunit. Because of this feature, PPMs encode multiple isoforms to achieve functional specificity (7). These enzymes depend on Mg^2+^ and Mn^2+^ cations to dephosphorylate proteins in eukaryotes. Generally, PP2C family members are abbreviated PTC (protein two C) enzymes. Presently, the human genome comprises 16 PP2C-encoding genes that undergo alternative splicing to produce 22 different isoforms, while *Arabidopsis thaliana* and *Oryza sativa* phosphatomes contain 80 and 78 PP2C-encoding genes, respectively (9–11). These proteins are involved in various biological processes and signal transduction during the growth and development of human and plants. The rice blast fungus phosphatome encodes 6 putative type 2C protein phosphatases, namely, MoPtc1, MoPtc2, MoPtc5-7 and MoPtc8. However, their precise roles in the pathophysiology of the fungus remains elusive.

The phosphatome of budding yeast comprises of 7 PP2C encoding genes namely (ScPtc1-ScPtc7) and ScPtc1 is best characterized. Previous findings have shown that ScPtc1-ScPtc4 play important roles in the deactivation of the osmosensing High Osmolarity Glycerol (HOG) pathway by dephosphorylating serine or threonine residues of the MAPK-HOG kinase as well as Pbs2 under acute osmolarity (12, 13). According to the results of a yeast two-hybrid assay, Ptc1 interacts with Pbs2 through the adaptor protein Nbp2 in *Saccharomyces cerevisiae* to regulate the HOG pathway (14). Similarly, Ptc1 dephosphorylates a series of MAPK protein kinases, including PKC, BCK1, MKK1/2, and SLT2, during cell wall stress in budding yeast. Ptc1 deletion mutants are sensitive to cell wall-compromising compounds and displayed increased phosphorylation levels of the Slt2 protein kinase (13, 15). (16). Among other functions, Ptc1 is also involved in the sporulation, inheritance and distribution of organelles, such as vacuoles, mitochondria and the cortical endoplasmic reticulum, tRNA splicing, cation homeostasis and the TOR pathway in budding yeast (17–20). Ptc1-disrupted cells are hypersensitive to LiCl, ZnCl, CaCl and rapamycin in *S. cerevisiae* (21). Moreover, Ptc2 and Ptc3 regulate the cell cycle by dephosphorylating the Thr-169 residue of the Cdc28 kinase (22). The regulation of the MAPK pathway in response to osmotic stress, oxidative stress and nutrient starvation is mediated by Ptc1 and Ptc3 in *S. pombe*. This is achieved through the direct dephosphorylation of SPC1-MAPK and the downstream transcription factor Atf1. Ptc1 has also been identified as a negative regulator of the cell wall integrity pathway in *S. pombe* and filamentous fungi such as *Neurospora crassa, Fusarium graminearum* and *Botrytis cinerea*(23–26). Recent findings have confirmed that XB15, a gene encoding a type 2C protein phosphatase, is involved in the negative regulation of disease resistance and the cell death response pathway in rice. The overexpression of XB15 resulted in increased resistance to bacterial pathogens, while the inactivation of XB15 had a contrasting effect on rice immunity (27). A search through genome for type 2C protein phosphatases in filamentous fungi such as *N. crassa, F. graminearum, B. cinerea* and *Aspergillus spp*. revealed that PP2C encoding genes are widely distributed in all filamentous fungi and six are harboured in *M. oryzae* genome suggesting that PP2C may be involved in major biological and cellular processes as in budding and fission yeast (8). However, their functions in filamentous fungi are poorly understood

The hemibiotrophic ascomycete *Magnaporthe oryzae* causes economically devastating blast disease in rice, wheat and other domesticated grass families (28). The annual losses resulting from rice blast disease are between 10-30%, posing a threat to global food security (29). The causal agent, *M. oryzae*, is considered a good model for understanding the molecular mechanisms underlying pathogen-host interactions (30). The disease cycle is initiated after viable conidia dispersed by wind or water droplets land on a susceptible host. The subsequent secretion of sticky mucilage from the apex of the conidium occurs to tightly adhere the conidium to the host surface.

Then, a polarized germ tube of approximately 15-30 μm emerges from the tip of the spore upon hydration by dew or water droplets (31, 32). The conidium then terminates germination and differentiates into a dome-shaped cell known as the appressorium in response to physical and chemical cues perceived on the host surface. This infective dome-shaped virulent cell structure is separated from the germ tube by a septum (33, 34). Appressorium development is regulated by multiple signalling pathways involving cAMP and MAPK in *M. oryzae* (35, 36). The highly conserved PMK1-MAPK, a functional homologue of *S.cerevisiae* Fuss/kss1, is required for the invasive growth and pathogenicity of *M. oryzae* and other appressorium-forming pathogens, such as *Colletotrichum lagenarium* and *Bipolaris oryzae* (37, 38). Mps1-MAPK is crucial for cell wall integrity, appressorium penetration and invasive growth in *M. oryzae*, and the MAPK-Osm1 pathway is activated in response to hyperosmotic stress, resulting in the accumulation of arabitol and glycerol in rice blast fungus (39, 40). Despite these physiological and pathological defects, the regulation of these MAPK signalling pathways remains elusive in the causative agent of rice blast. In this study we first evaluated the transcription expression level of five type 2c protein phosphatase encoding genes and found that these genes are differentially expressed during pathogen-host interaction. Among the five genes MoPtc1, MoPtc2 and MoPtc7 were significantly upregulated at appressorium development (12h) as compared to the other type 2c protein phosphatase encoding genes. MoPtc1 and MoPtc2 were further characterized and found to be necessary for vegetative growth, conidiation and regulating MAPK signalling pathways in *M. oryzae*.

## Results

### Identification of MoPtc1 and MoPtc2 *in M. oryzae*

Through protein blast search (BLASTP) using *S. cerevisiae* type2C protein phosphatases amino acid sequences as protein queries, we identified five single PP2C orthologs in the genome of *M. oryzae* with exception of *S. cerevisiae* Ptc3 and Ptc4. The new member Ptc8 is not encoded in *S. cerevisiae* genome hence was identified in *M. oryzae* by protein blast search (BLASTP) using *C. albicans* CaPtc8 amino acid sequence. Among the five members MGG_05207 and MGG_01351 encode for MoPtc1 and MoPtc2 respectively. The primary structures and percentage identities with *S. cerevisiae* and *C. albicans* counterparts are described in Table 1.

**Table 1.**
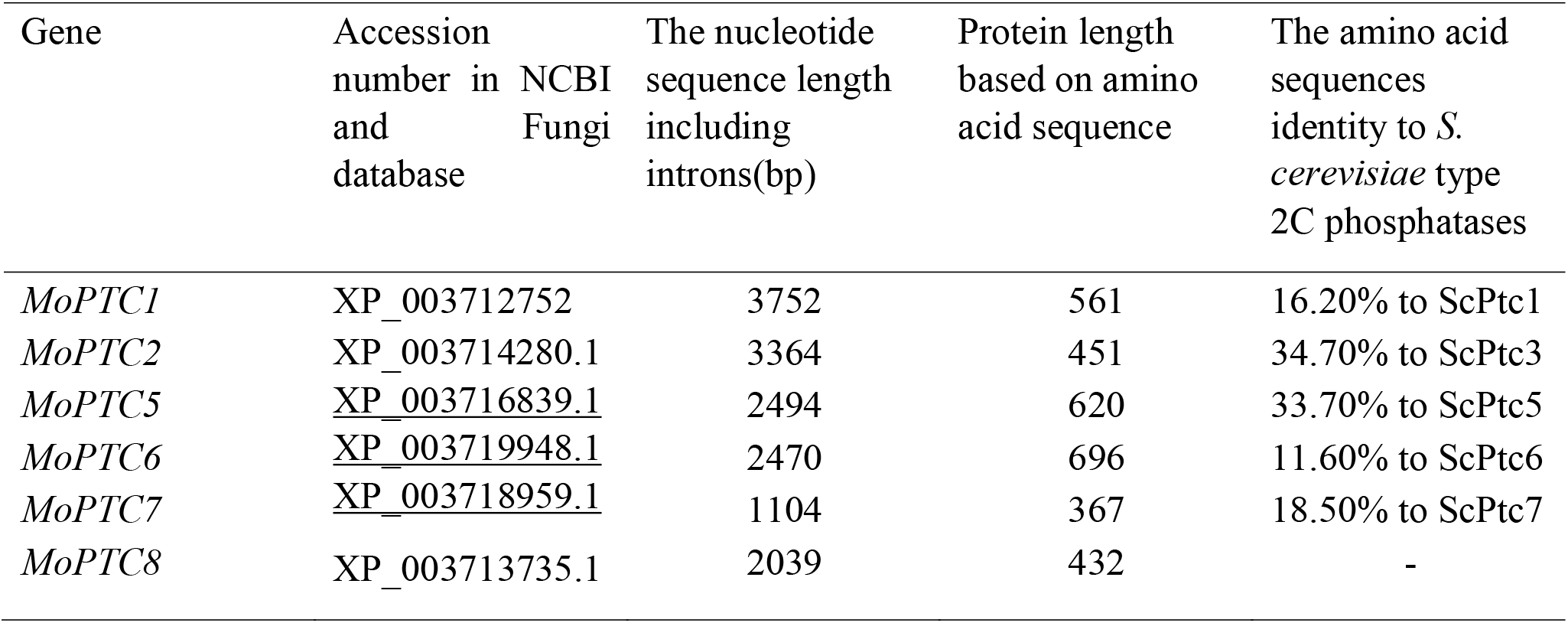
Primary structure analysis of six type 2C protein phosphatases in *M. oryzae*.

| Gene | Accession number in NCBI and Fungi database | The nucleotide sequence length including introns(bp) | Protein length based on amino acid sequence | The amino acid sequences identity to *S. cerevisiae* type 2C phosphatases |
| --- | --- | --- | --- | --- |
| *MoPTC1* | XP_003712752 | 3752 | 561 | 16.20% to ScPtc1 |
| *MoPTC2* | XP_003714280.1 | 3364 | 451 | 34.70% to ScPtc3 |
| *MoPTC5* | <u>XP_003716839.1</u> | 2494 | 620 | 33.70% to ScPtc5 |
| *MoPTC6* | <u>XP_003719948.1</u> | 2470 | 696 | 11.60% to ScPtc6 |
| *MoPTC7* | <u>XP_003718959.1</u> | 1104 | 367 | 18.50% to ScPtc7 |
| *MoPTC8* | XP_003713735.1 | 2039 | 432 | - |

Analysis of the domain architecture by SMART database (http://smart.embl-heidelberg.de/) revealed that MoPtc1 (MGG_05207) contains a PP2Cc (Serine/threonine phosphatase, family 2C, catalytic domain) at positions 141-478 and PP2C_SIG (Sigma factor PP2C-like phosphatase) domain at position 190-480 amino acid respectively (Fig. 1). The PP2Cc catalytic domain contains the typical phosphatase active site with –ED-DGH (A/G)-GD-GD-DG-DN-signature motif sequence, whereas the PP2C_SIG domain harbours the catalytic subunit of -DGH (A/G)-GD-GD-DG-DN-signature motif. MoPtc2, on the other hand, is a 461 amino acid long protein that contains PP2Cc domain at position 13-293 and PP2C_SIG domain at position 39-295 respectively (Fig. 1).

**Figure 1.**
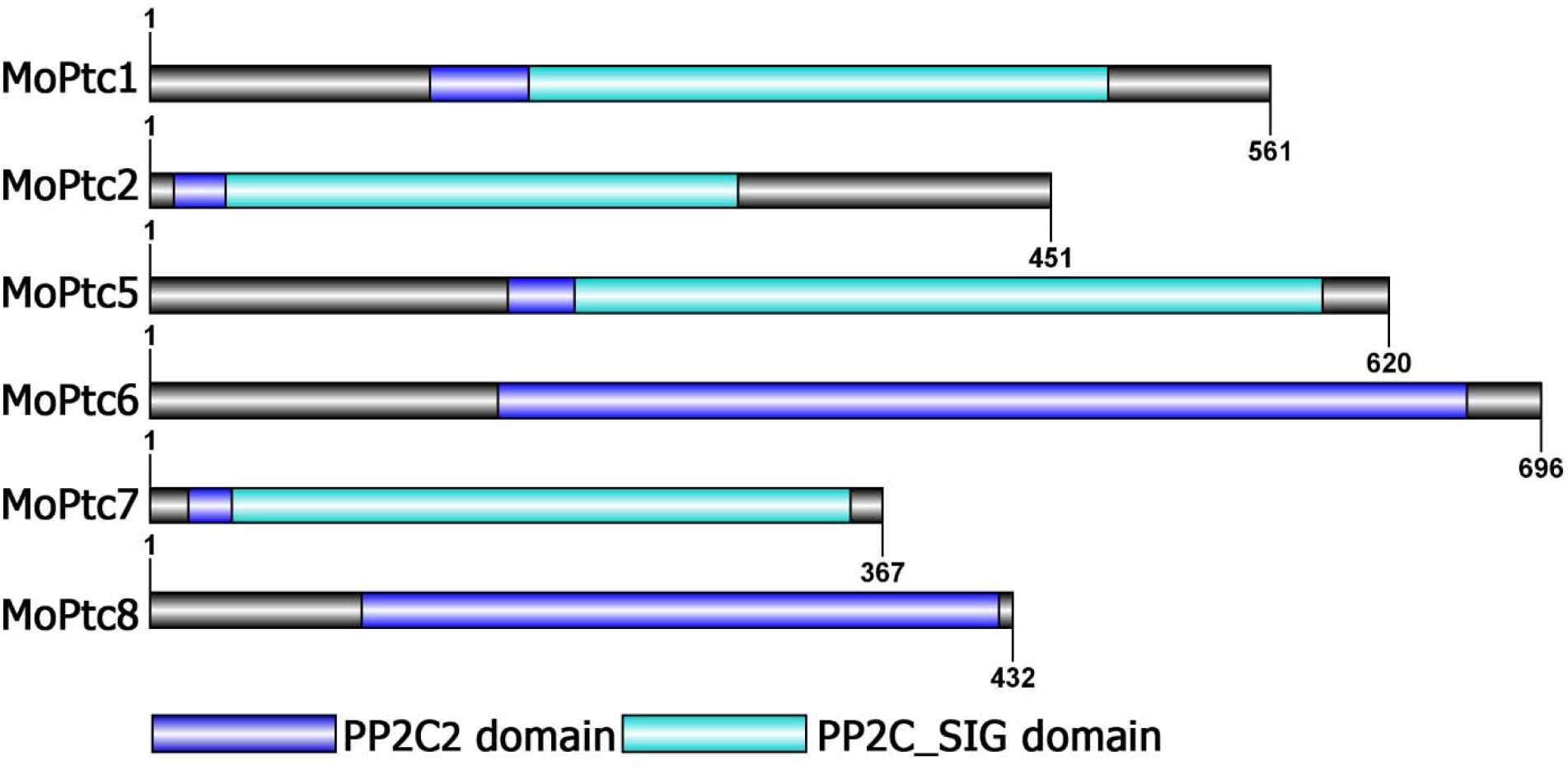
Domain architecture of type 2C protein phosphatases in *Magnaporthe oryzae*. The image was generated by using IBS software of version 1.0.3. The domains were predicted with SMART database.

### MoPtc1 and MoPtc2 are expressed during host-pathogen interactions as well as by multiple stress causing agents

Firstly, we conducted the quantitative real time PCR (qRT-PCR) to monitor the expression levels of type 2c protein phosphatases encoding genes during pathogen-host interactions. Three weeks old susceptible rice seedlings were sprayed with Guy11 spore suspensions and RNA was extracted from these infected plant tissues at different time intervals. From the results, MoPtc1 and MoPtc2 were expressed at different developmental stages of infectious cycle and showed significant expression at 12h, of post incubation (Fig.2A). These findings suggest that MoPtc1 and MoPtc2 may play important roles in appressorium development in *M. oryzae*.

**Figure 2.**
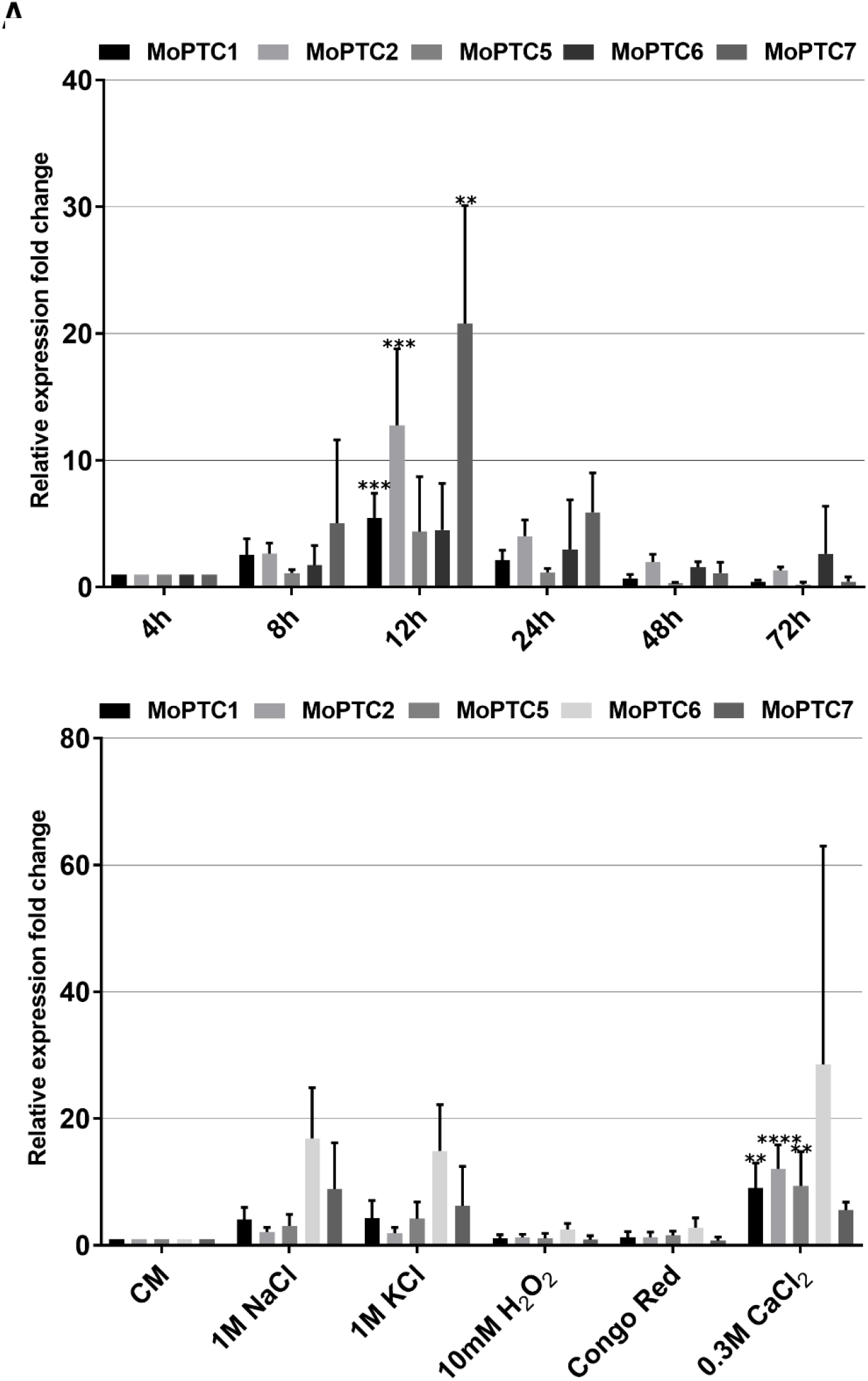
The Transcript expression profile of *Magnaporthe oryzae* type 2c protein phosphatase encoding genes. **(A)** In planta expression pattern of *MoPTC1, MoPTC2, MoPTC5, MoPTC6* and *MoPTC7, at* 4, 8, 12, 24, 48 and 72 hours of infection. **(B)** Transcript expression pattern of *MoPTC1, MoPTC2, MoPTC5, MoPTC6* and *MoPTC7* in different stress causing agents. In these assay, β-tubulin was used as reference gene. The error bars represent the mean standard error from three independent replicates. The double asterisks indicate adjusted P value of 0.0013, triple asterisks denotes adjusted P value of 0.0007 and Quadruple asterisks shows adjusted P value of <0.0001 respectively.

Next, we quantified the transcript abundance of five type 2c protein phosphatase encoding in culture media containing different stress and found that MoPtc1, MoPtc2, MoPtc5, MoPtc6 and MoPtc7 are differentially expressed in osmotic (NaCl and KCl), and ionic (CaCl_2_) stressors with the highest transcript expression upon exposure to ionic stress CaCl_2_ (Fig. 2B). These results support the idea that MoPtc1 and MoPtc2 are involved in regulating multiple stress responses in *M. oryzae*

### The deletion of MoPtc1 and MoPtc2 has significant effect towards cell wall stress agent

To reveal the functions of MoPtc1 and MoPtc2 in *M. oryzae*, the gene deletion construct was amplified by double joint PCR and transformed into protoplasts of the wild-type strain Guy11. The obtained transformants were screened by PCR, and the resulting mutants were confirmed by Southern blotting (Fig. S1). For each gene only one mutant was used for further analysis. The genetic complementation of Δ*Moptc1* and Δ*Moptc2* mutants was performed by transforming MoPtc1-GFP and MoPtc2-GFP with a native promoter into the deletion mutants.

We investigated the contribution of MoPtc1 and MoPtc2 in osmotic stress response and cell wall integrity of the rice blast fungus. Ten days post-incubation, both Δ*Moptc1* and Δ*Moptc2* showed insignificant change in sensitivity towards hyperosmotic stressors (NaCl, KCl and Sorbitol), oxidative stress agent (H_2_O_2_) and cell wall and cell membrane stress agents (CFW & SDS) respectively (Fig. 4B). However, the single deletion mutant and double deletion mutants MoPtc1 and MoPtc2 displayed a significant growth inhibition in media supplemented with Congo red which is a cell wall stress inducing agent (Fig. 3B). These results suggest that MoPtc1 and MoPtc2 play a crucial role in maintaining cell wall integrity in rice blast fungus (Fig. 3B).

**Figure 3.**
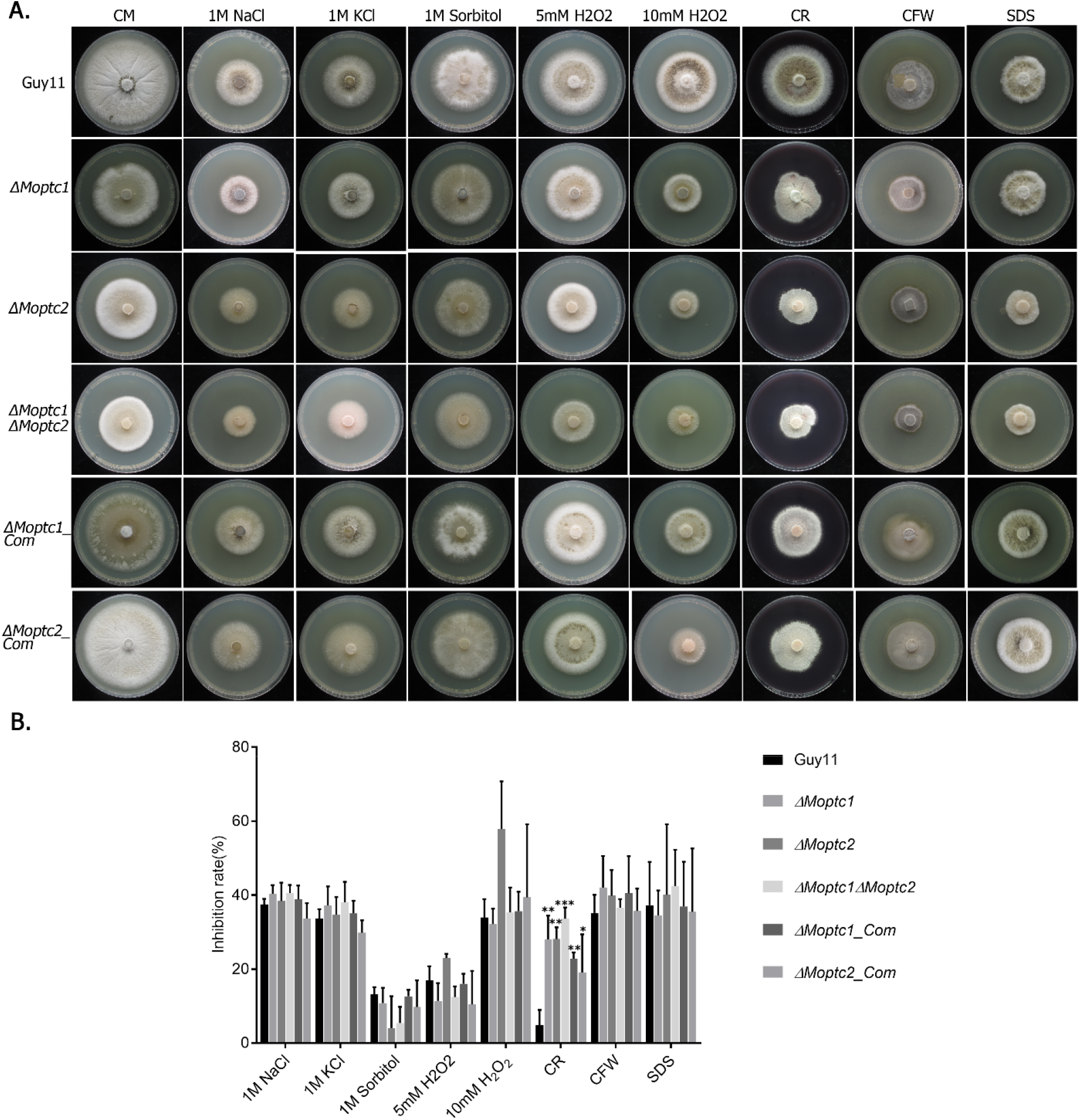
The deletion of MoPtc1 and MoPtc3 reduces tolerance to cell wall stress agent. **(A)** Growth inhibition assay of Guy11, Δ*MoPtc1*, Δ*MoPtc3*, Δ*MoPtc1*Δ*MoPtc3*, Δ*MoPtc1_com, and* Δ*MoPtc3_com* strains in the presence of osmotic, oxidative and cell wall stress. **(B)** Bar charts showing the inhibition difference between Guy11 and deletion strains in the presence of osmotic, oxidative and cell wall stressing agents. The error bars represent the mean standard error from three independent replicates (** shows adjusted P Value of 0.0011, *** indicates adjusted P value of 0.0002, and * represents adjusted P Value of 0.0319 respectively).

**Figure 4.**
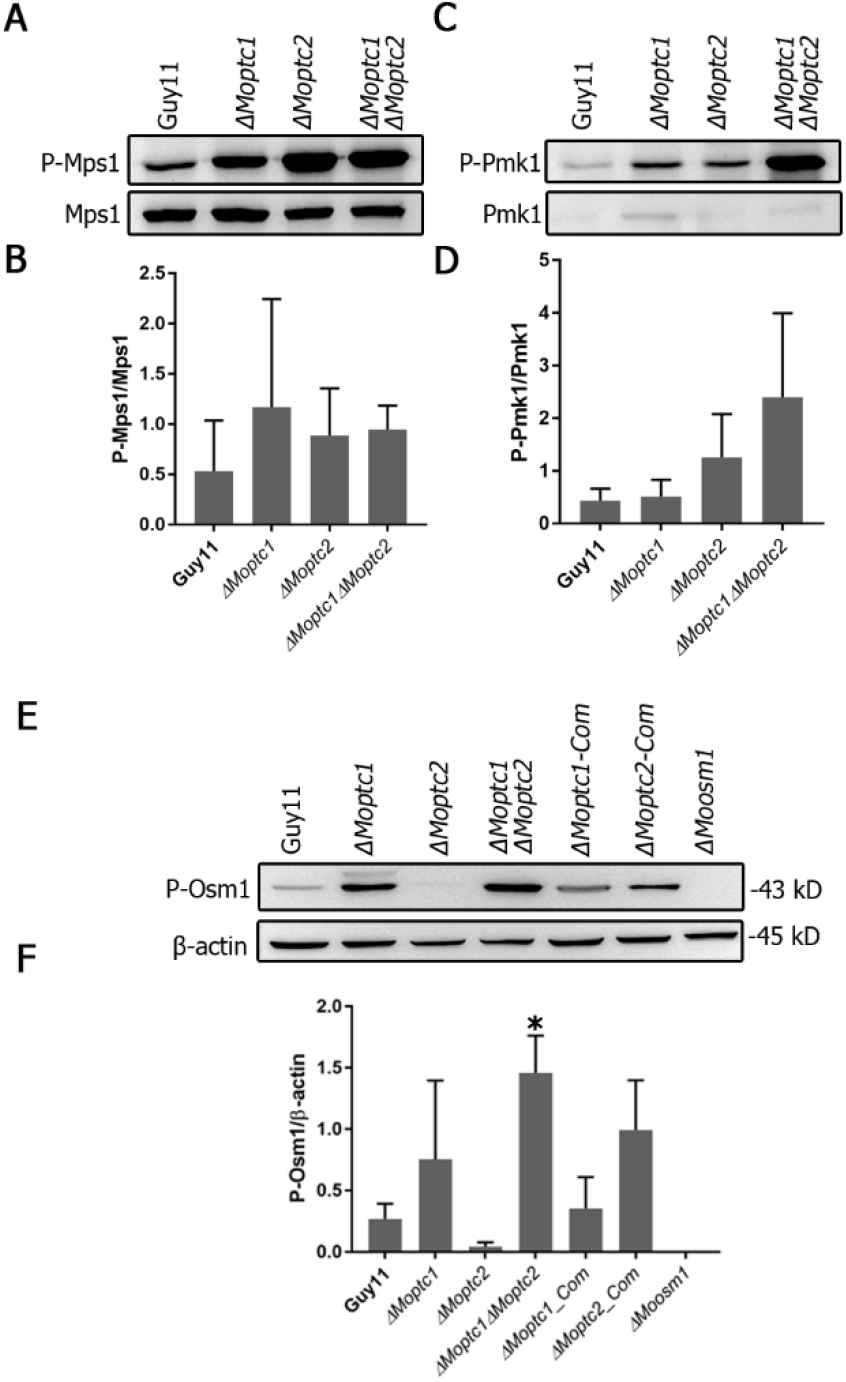
MoPtc1 and MoPtc3 regulate MAPK signaling cascades in *M. oryzae*. **(A)** Immunoblot assay showing the phosphorylation level of Mps1 in Guy11, Δ*Moptc1*, Δ*Moptc3* and Δ*Moptc1*Δ*Moptc3* strains. The strains were shaken in complete media (CM) and protein extracted for western blot analyses. Equal amounts of protein were loaded into each well and the phosphorylation level was detected using phospho-p44/42 MAPK (ERK1/2) (Thr202/Tyr204) (D13.14.4E) rabbit mAb and p44/42 MAPK (Erk1/2) antibodies respectively. **(B)** The band intensity ratio for Phosphorylated Mps1 (P-Mps1) compared with the control (Mps1). These ratios were calculated by dividing p-Mps1/Mps1. **(C)** Western blot assay showing the Pmk1 phosphorylation level in Guy11, Δ*Moptc1, and* Δ*Moptc3* and Δ*Moptc1*Δ*Moptc3* strains. The strains were shaken in complete media (CM) and protein extracted for western blot analyses. Equal amounts of protein were loaded into each well and the phosphorylation level was detected using phospho-p44/42 MAPK (ERK1/2) (Thr202/Tyr204) (D13.14.4E) rabbit mAb and p44/42 MAPK (Erk1/2) antibodies respectively. **(D)** The band intensity ratio for Phosphorylated Pmk1 (P-Pmk1) compared with the control (Pmk1). These ratios were calculated by dividing p-Pmk1/Pmk1. **(E)** Western blot image quantifying the Osm1-MAPK phosphorylation in Guy11 and Δ*MoPtc1*, Δ*MoPtc3* Δ*MoPtc1*Δ*MoPtc3* strains. The strains were shaken in complete media (CM) and protein extracted for western blot analyses. The phosphorylation level of MoOsm1 in all strains was detected using phosphop38 MAPK (Thr180/Tyr182) (D3F9) XP rabbit mAb while β-actin was detected using actin (2P2) mouse mAb. Equal amounts of protein were loaded into each well. **(F)** The band intensity ratio between phosphorylated Osm1 (P-Osm1) and β-actin. The single asterisk (*) represent statistical significance with adjusted P Value of 0.0338.

### MoPtc1 and MoPtc2 play overlapping roles in regulating MAPK signalling pathways in *M. oryzae*

To investigate the role of MoPtc1 and MoPtc2 in the regulation of the MAPK signalling pathway, we assayed the phosphorylation level of MoPmk1, MoMps1 and MoOsm1 by Western blot. The phosphorylation levels of Mps1 and Pmk1were increased in both ΔMoptc1 and Δ*Moptc2* single deletion mutants and Δ*Moptc1*Δ*Moptc2* double deletion strains compared to Guy11 (Fig. 4A&4B). For Osm1, total protein was extracted from the mycelia of Guy11 and Δ*Moptc1*, Δ*MoPtc2*, Δ*Moptc1*Δ*MoPtc2*, Δ*Moptc1_Com*, and Δ*MoPtc2_Com* strains. A significant increase in Osm1 phosphorylation was detected in Δ*Moptc1*, and Δ*Moptc1*Δ*Moptc2* mutants while MoPtc2 was decreased in Osm1 phosphorylation levels (Fig. 4E). These results strongly suggest that MoPtc1 and MoPtc2 played synergistic roles in dephosphorylating Pmk1-MAPK, Mps1-MAPK and Osm1-MAPK in *M. oryzae*. In addition, MoPtc1 is a negative regulator of Osm1 during hypoosmotic stress while MoPtc2 positively regulates this pathway.

### MoPtc1 interacted with Pmk1 and MoOsm1 via the adaptor protein MoNbp2 both in vitro and in vivo

We assayed for possible interaction between MoPtc1 or MoPtc2 with Pmk1, Mps1 or Osm1 in *M. oryzae* via yeast two hybrid assays. The results showed a strong interaction between MoPtc1 and the adaptor protein MoNbp2 (Fig. 5A) and between MoNbp2 and Pmk1 (Fig. 5B), and between MoNbp2 and MoOsm1 (Fig. 5C) in *M. oryzae*. These interaction was further confirmed by co-immunoprecipitation (Co-ip) assay in vivo. First we co-transformed Nbp2-Flag into the protoplast of Guy11-GFP as our control experiment and was able to detect a 64kd band size of Nbp2 using anti-Flag antibody as well as GFP band size of 27kD using anti-GFP antibody (Fig. 5D). Secondly we co-transformed Nbp2-Flag and Ptc1-GFP constructs into Guy11 protoplast and was able to detect an 88kD band size corresponding to Ptc1-GFP and 64kD band size for Nbp2-Flag in total proteins and GFP magnetic beads. (Fig. 5D). Thirdly we co-transformed Pmk1-GFP and Nbp2-Flag constructs into Guy11 protoplast and were able to detect both the Pmk1-GFP (66kD) band and Nbp2-Flag (64kD) band in the total proteins and co-immunoprecipitation lysates (Fig. 5E). Finally we co-transformed Osm1-GFP and Nbp2-Flag constructs into Guy11 protoplast and found that the anti-Flag antibody was able to detect specifically a clear band of 64kD while anti-GFP antibody was able to detect a clear band of 66kD (Fig. 5F). Together these results demonstrate that MoPtc1 interacts with Pmk1 and Osm1 via the adaptor protein in M. oryzae.

**Figure 5.**
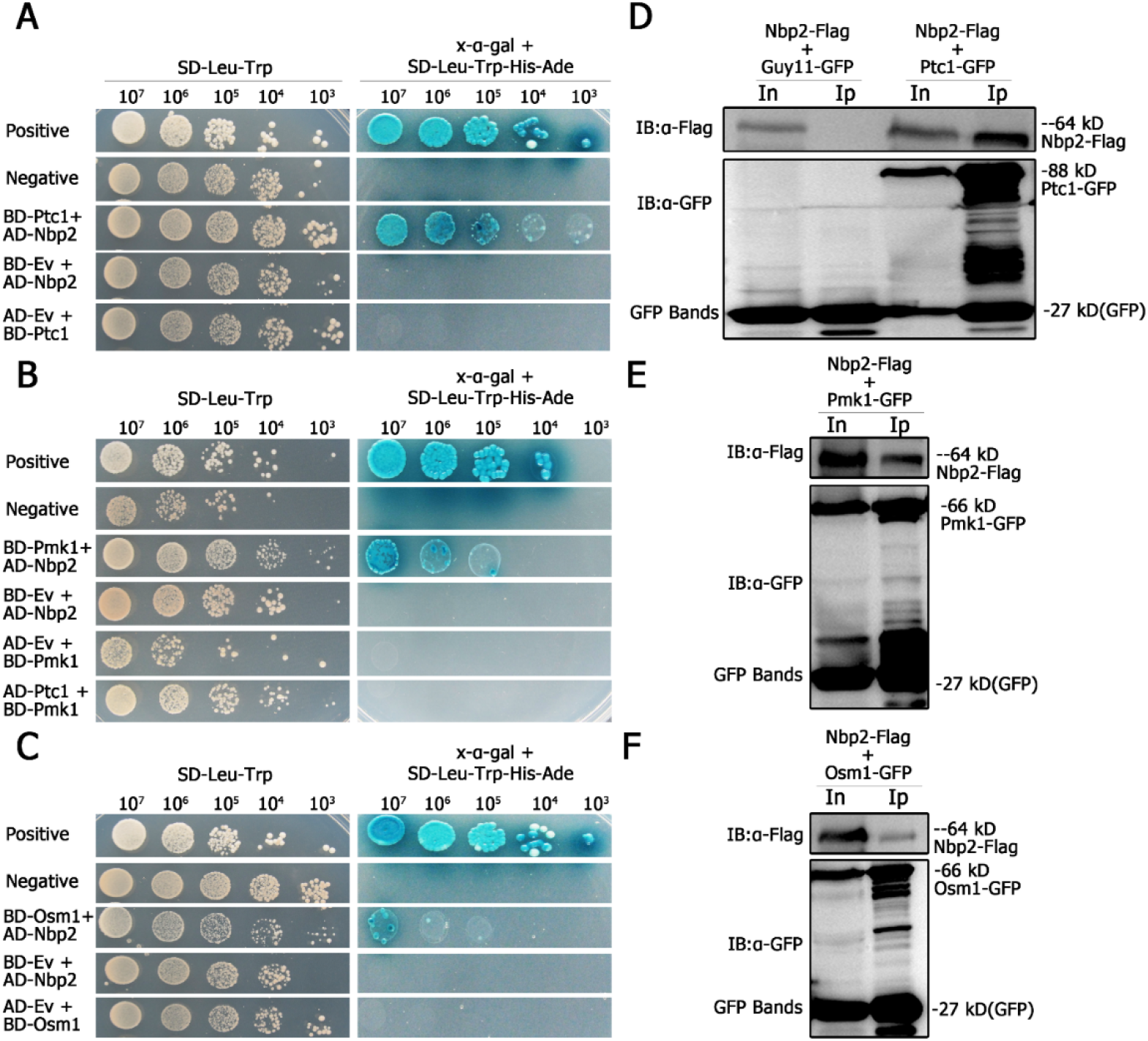
MoPtc1 interacts with Pmk1 and MoOsm1 in *M. oryzae*. Yeast two hybrid showing interaction between **(A)** MoPtc1 and MoNbp2 and between **(B)** MoNbp2 and MoPmk1 and between **(C)** MoNbp1 and MoOsm1 in rice blast fungus. The pGBKT7-53/pGADT7-T and pGBKT7-Lam/pGADT7-T vectors were used as positive and negative controls respectively. The Cotransformation of prey and bait constructs into yeast strain resulted in transcription activation of reporter genes and growth on the selective medium -Leu/-Trp/-His/-Ade. The yeast colonies turned blue after hydrolysis of X-gal and secretion of β-galactosidase (LacZ) in the selective medium. CoImmunoprecipitation assay confirming the interaction between **(D)** MoPtc1 and MoNbp2 and between **(E)** MoNbp2 and MoPmk1 and between **(F)** MoNbp1 and MoOsm1 in rice blast fungus.

### Type 2c protein phosphatases are localized in different cellular compartments in *M. oryzae*

To determine the subcellular localization of type 2c protein phosphatases we first co-transformed MoPtc1-GFP and MoPtc2-Mcherry with a constitutive promoter in Guy11 protoplasts and found that MoPtc1 colocalized with MoPtc2 in the cytoplasm in mycelium (Fig. 6A) and invasive hypha (Fig. 6K). We also checked for the localization of MoPtc1, MoPtc2, MoPtc5 and MoPtc6 by transforming the MoPtc1-GFP, MoPtc2-GFP, MoPtc5-GFP and MoPtc6-GFP under a constitutive promoter into the protoplast of Guy11 and found that MoPtc1 is localised in the nucleus and cytoplasm in both conidia (Fig. 6C) and appressorium (Fig. 6G) stages of development while MoPtc2 was localised in the cytoplasm (Fig. 6D & 6H). MoPtc5 on the other was localised in mitochondria in all stages of development (Fig. 6B, 6E, 6I & 6L) while MoPtc6 was localised in cytoplasm in conidia and appressorium stages (Fig. 6F & 6J). Taken together our results demonstrate that type 2c protein phosphatases are localized at different cellular compartments in the life cycle of Magnaporthe oryzae.

**Figure 6.**
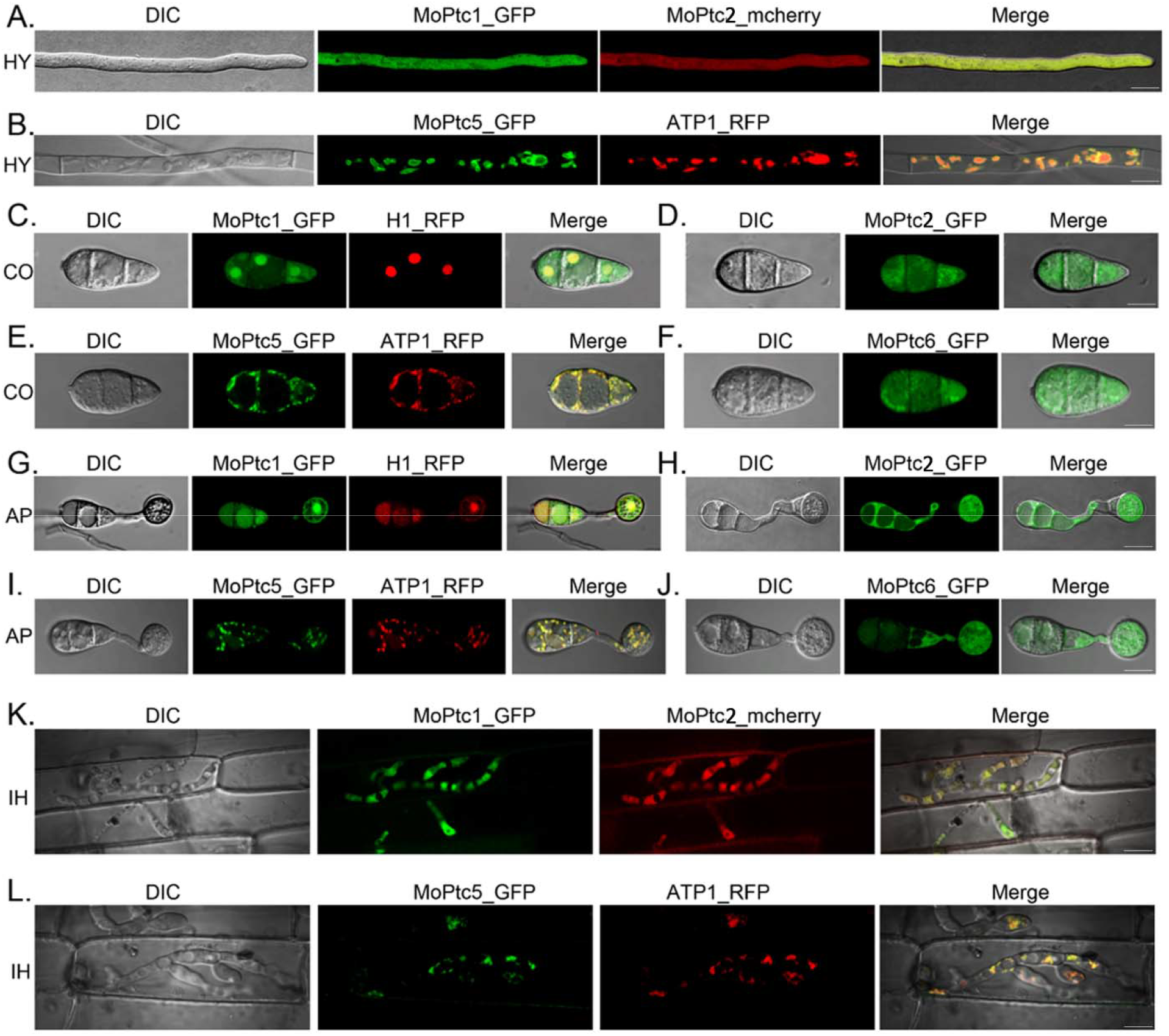
Subcellular localization of type 2c protein phosphatases in M. oryzae. **(A)** Colocalization of MoPtc1 and MoPtc3 in hyphal (HY) stages of development **(B)** Localization pattern of MoPtc5 in hyphae. **(C-F)** Localization of MoPtc1, MoPtc3, MoPtc5 & MoPtc6 in conidia (CO). **(G-J)** The localization of MoPtc1, MoPtc3, MoPtc5 & MoPtc6 in appressorium (AP) development. **(K)** Colocalization of MoPtc1 and MoPtc3 in invasive hypha (IH). **(L)** Localization of MoPtc5 in invasive hypha. The live cell images were visualized by Nikon Air Laser confocal microscopy (scale bar = 20 μm).

## Discussion

Type 2C protein phosphatases regulate a wide range of signaling pathways and biological processes in eukaryotes. However their functions in the MAPK signalling and pathophysiology of *M. oryzae* remains elusive. In this study we set out to determine the roles of MoPtc1 and MoPtc2 in MAPK signalling and pathogenesis of *M. oryzae*. Firstly, we identified the type 2c protein phosphatase homologues (MoPtc1, MoPtc2, MoPtc5-7) in *M. oryzae* using a protein blast (BLAST-P) program of fungi and oomycete with *Saccharomyces cerevisiae* as query sequence and found that *M. oryzae* contains a single copy of MoPtc1, MoPtc2, and MoPtc5-7 respectively. Next, we predicted the domains of these proteins using SMART data base and found that MoPtc1, MoPtc2, MoPtc5 and MoPtc7 have PP2C and PP2C_SIG domains while MoPtc6 and MoPtc8 have PP2C domain only. These results may suggest that PP2C domain is conserved among the type 2c protein phosphatases while PP2C_SIG domain is not.

Through transcription analysis, MoPtc1 and MoPtc2 were significantly and differentially upregulated at 12h of infection. These results show that both genes may be involved in appressorium mediated infection of plant tissues. Additionally MoPtc1 and MoPtc2 were differentially induced by different stress causing agents and showed a significant increase in the presence of 0.3M CaCl_2_.These results suggest that MoPtc1 and MoPtc2 may be important in ionic stress responses in rice blast fungus.

In plant pathogenic fungi, Slt2 pathway is crucial for asexual reproduction, cell wall integrity and pathogenesis. Not only was the Mps1 null mutants in *M. oryzae* reduced in aerial hypha growth but also in conidiation and tolerance to cell wall lysis enzymes^(39)^. Different from Pmk1 mutants, the *Mps1* mutants in rice blast fungus formed appressorium but could not repolarize leading to a complete loss of pathogenicity^(39)^. Additionally, *M. oryzae* contains novel receptors involved in cell wall stress signal perception because among all the cell surface receptors encoded by budding yeast only Wsc1 displayed a limited homology to one gene in *M. oryzae*^(47)^The Pmk1 pathway plays a crucial role in infection related morphogenesis^(48)^ while the Mps1 plays an important role in cell wall integrity in rice blast fungus. By immunoblot analysis we found that the phosphorylation level of MoPmk1-MAPK, MoMps1-MAPK was increased in both Δ*Moptc1 and* Δ*Moptc2* single deletion mutants and Δ*Moptc1*Δ*Moptc2* double deletion mutants suggesting that MoPtc1 and MoPtc2 play synergistic roles in regulating negatively the MoPmk1 and MoMps1 pathways. Our results are similar to those in budding yeast and other filamentous fungi such as *M. oryzae, N. crassa, F. graminearum* and *B. cinerea* where Ptc1 is involved in regulating negatively the cell wall integrity pathway(23–26).

The High osmolarity glycerol (HOG) pathway plays a key role in adaptive responses to osmotic, heat, and oxidative stress^(49)^. This pathway also mediates sensitivity to certain fungicides like phenylpyrroles and dicarboximides in filamentous fungi and budding yeast^(50)^. High concentrations of hydrogen peroxide have also been shown to activate the HOG pathway in *S. cerevisiae*^(51, 52)^, however the mechanisms involving stress signal sensing and relay is yet to be elucidated at molecular level. Deletion of MoPtc1 resulted in a dramatic increase in the phosphorylation of MoOsm1, while the phosphorylation level of MoOsm1 was decreased or lower in Δ*Moptc2* deletion mutants. These findings suggest that MoPtc1 exerts a negative effect on the HOG pathway while MoPtc2 exerts a positive effect during hypoosmotic stress in *M. oryzae*. In addition, MoPtc1 may play a crucial role in maintaining low basal MoOsm1 phosphorylation levels under normal conditions. Taken together, MoPtc1 and MoPtc2 play synergistic roles in dephosphorylating MAPK signalling cascades in *M. oryzae*.

It has previously been shown in budding yeast that MoPtc1 is recruited to MoPbs2 and Hog1 via adaptor protein MoNbp2 to regulate the High osmolarity glycerol (HOG) pathway during high osmotic stress ^(53)^. Ptc1 interacted with Nbp2, an adaptor protein in *S. cerevisae* to dephosphorylate MAPK protein kinase Pbs2 under acute osmolarity^(53)^. Pbs2 is a scaffolding protein of STE11, HOG1, and SSK2/22 and STE11 of MAPK pathway. Similar to budding yeast, MoPtc1 strongly interacts with MoNbp2; however, MoNbp2 interacts with MoOsm1, MoPmk1 and not MoPbs2 in *M. oryzae*. This is the first report in our knowledge about the regulation of MAPK signaling pathways through adaptor proteins in plant pathogenic fungi.

In budding yeast, Ptc1 deletion mutants were sensitive to cell wall antagonists as caffeine, caspofungin or Calcoflour white and to conditions that activate the CWI pathways such as alkaline pH^(54)^. *M. oryzae* senses and responds to cell wall stresses through CWI pathway during development and pathogenicity (39, 55). The deletion of MoPtc1 and MoPtc2 significantly compromised cell wall integrity in *M. oryzae*. MoPtc1 and MoPtc2 are negative regulators of the Mps1 pathway in *M. oryzae* and that the sensitivity of Δ*Moptc1 and* Δ*Moptc1*Δ*Moptc2* mutants towards agents that induce cell wall stress is due to the hyperactivation of the cell wall integrity pathway. Collectively, our results suggest that Ptc1 plays crucial role in maintaining cell wall integrity through dephosphorylation of Mps1 in *M. oryzae*

In conclusion, the type 2c protein phosphatases MoPtc1 and MoPtc2 play synergistic roles in regulating MAPK signalling pathways in *M. oryzae* as shown in the proposed model (Fig. 7). Understanding the regulatory mechanisms of MAPK signalling pathways could lead to the discovery of new anti-blast agents that will prevent the spread of rice blast disease.

**Figure 7.**
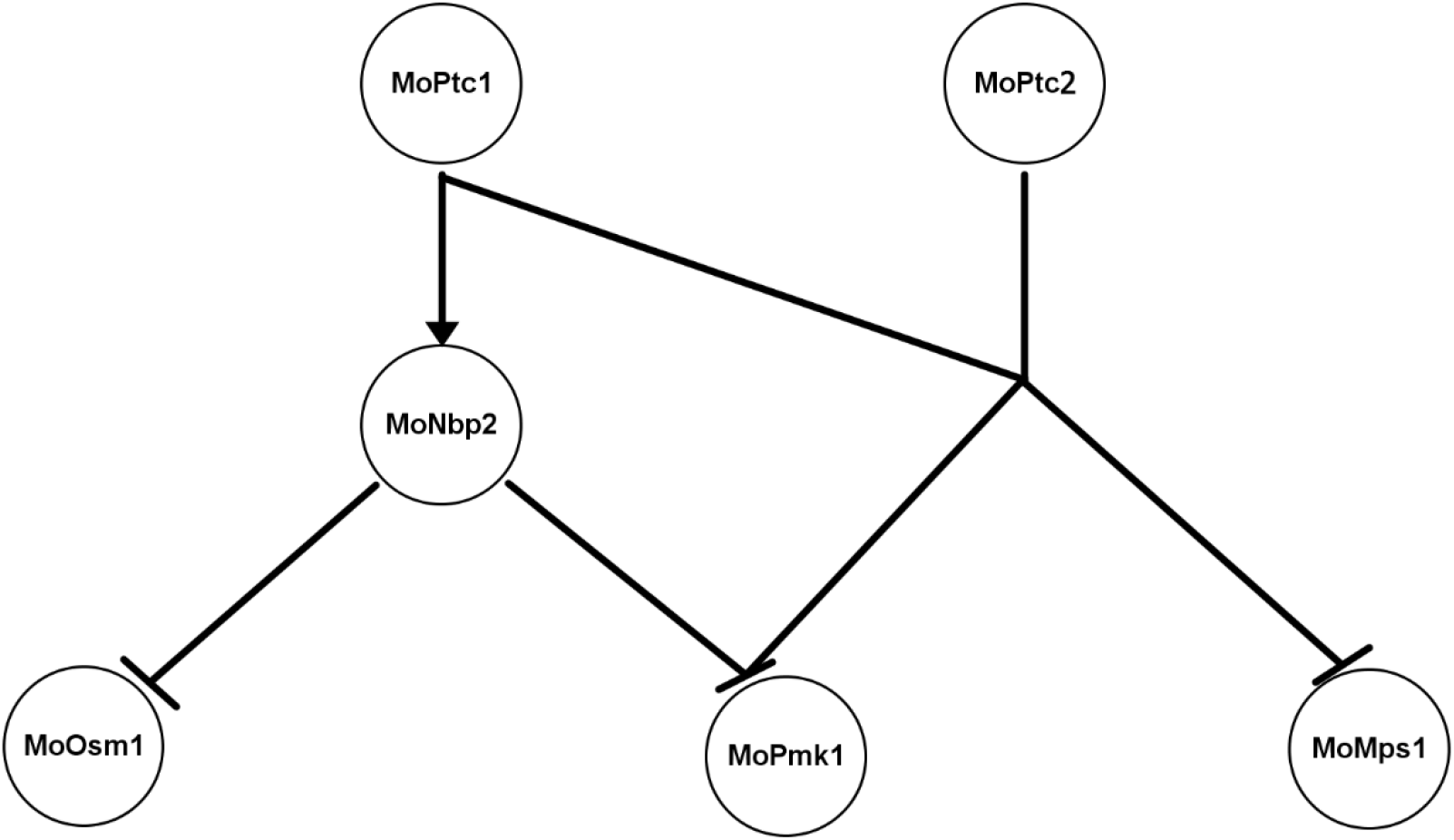
A proposed model showing regulation of MAPK signaling pathways by MoPtc1 and MoPtc2 in *Magnaporthe oryzae*. In this model MoPtc1 negatively regulates MoOsm1, MoPmk1 and MoMps1, while MoPtc2 negatively regulates two MAPK proteins namely MoPmk1 and MoMps1 in *M. oryzae*. However, MoPmk1 and MoOsm1 are negatively regulated by MoPtc1 via adaptor protein known as MoNbp2 in *M. oryzae*.

## Materials and Methods

### Fungal strains and culture conditions

*M.oryzae* Guy 11 was used as wild type strain in this study. Culturing of strains was done on complete medium (CM containing yeast extract-0.6%, casamino acid-0.6%, sucrose-1%, agar-2%). For cell wall sensitivity test, the strains were grown on complete solid media containing 200μg/mL Congo red, 0.01% SDS and 200μg/ml of CFW. The strains were incubated at 27□, and the colony diameters measured at 5, 7 and 10 days respectively as described in (56). These experiments were done at least three times with three repeats. For osmotic and oxidative stress assay, a plug of mycelia was inoculated on complete medium containing 1M NaCl, 1M KCl, 1M Sorbitol, 5mM H_2_O_2_ and 10mM H_2_O_2_ and incubated at 27 □ for 10 days.

### Generation of MoPtc1 and MoPtc2 deletion strains in *M. oryzae*

The split marker approach was used to generate the target gene replacement constructs of *MoPTC1* and *MoPtc2* in *M. oryzae*. The flanking sequences were amplified with the primer pairs MoPtc2-AF/AR and MoPtc2-BF/BR respectively (Table1.1). These fragments were then ligated to HY and YG split parts of Hygromycin resistant gene cassette by using gene splicing by overlap extension (SOE PCR). The primer used are also listed in the table below (Table1.1). The Polyethylene glycol (PEG) mediated Protoplast transformation was done in Guy 11 background strains as described (57, 58). The positive transformants were confirmed and verified using PCR and Southern blotting methods. To generate Δ*Moptc1*Δ*Moptc2* double deletion constructs, 1 kb and 0.9 kb flanking sequences were amplified with the primer pairs MoPtc2-AF/AR and MoPtc2-BF/BR. These fragments were then ligated with split parts of Neomycin resistant gene Open Reading Frame (NE and EO) by using gene splicing by overlap extension PCR (SOE-PCR). This construct was then transformed into Δ*Moptc1 mutant* protoplast and resulting transformants with G418 resistance were screened by PCR and verified by Southern blotting. Primers used for generation of these mutants are listed in Table 1.1.

### Quantitative real time PCR (qRT-PCR) and RT-PCR assays

The strains were shaken in liquid complete media at 130rpm, 28□ for 96 hours. Total RNA was extracted from the mycelia using Eastep™ Total RNA Extraction Kit (Promega). Complementary DNA (cDNA) was synthesized using PrimeScript™ RT reagent kit with gDNA Eraser (Perfect Real Time). quantitative real time PCR (qRT-PCR) was performed using 10μL reaction mixture containing 5μL SYBR® Premix Ex. Taq™, 3.4 μL DNase/RNase free water, 1 μL 10xcDNA and 0.3 μL of Forward and Reverse primers as recommended by Promega Corporation Super Real Premix. The qRT-PCR data was produced by Eppendorf Realplex2 master cycler (Eppendorf AG 223341, Hamburg) and analysis done using delta delta-CT (2□-□ΔΔCT) method as described by(59, 60). β-tubulin was used as reference gene and primers were designed using Beacon designer Software version 8 as listed in Table 1.1.

### Construction of MoPtc1 and MoPtc2 complementation vector with GFP fusion protein

MoPtc1-GFP and MoPtc2-GFP vectors were constructed by amplifying 3.5 Kb and 3.9 Kb full length ORF of MoPtc1 and MoPtc2 with their respective native promoter regions. The resulting PCR products were cloned into *Kpn*I / *Hin*dIII site of pKNTG plasmid having GFP expression sequence at the C-terminal. (Primers used are listed in Table 1.1). The fusion construct was sequenced to validate the orientation of the inserted fragment and transformed intoΔ*Moptc1 and* Δ*Moptc2* protoplast respectively. The resulting transformants with G418 resistance were screened by PCR method with primer pairs MoPTC1 OF -GFP/R and MoPtc2 OF-GFP/R respectively. The GFP signals were screened under Olympus BX51 microscopy (Olympus, Japan) and the pictures were captured by Nikon A1R laser scanning confocal microscopy system (Nikon, Japan)

### Immunoblotting assays

The strains were shaken in complete media (CM) for 72 hours and mycelia were ground to fine powder in liquid nitrogen and protein extracted and resuspended in 1mL protein lysis buffer (10mM Tris-HCl pH-7.5, 150mM NaCl, 0.5mM EDTA, 0.5% Nonidet P-40) in the presence of 5μL protease inhibitor cocktail (Sigma-Aldrich’s. Louis, USA) and 1mM PMSF. The total protein was separated on 12.5% SDS-PAGE gels and transferred to nitrocellulose membranes (Amersham, Piscataway, NJ, USA) for western blot analyses. The membranes were incubated with phospho-p38 MAPK (Thr180/Tyr182) (D3F9) XP Rabbit mAb, phospho-p44/42 MAPK (ERK1/2) (Thr202/Tyr204) (D13.14.4E) Rabbit mAb and p44/42 MAPK (Erk1/2) antibodies (Cell Signaling Technology, Beverly, MA, USA) and affinity purified goat anti-rabbit & mouse antibody conjugated to horseradish peroxidase (Abmart, Shanghai, China).The ECL kit (Amersham Biosciences, Germany)was used to detect the chemiluminescent signals of the specific protein bands(61). The protein samples were also detected with actin (2P2) mouse mAb as reference (Abmart, Shanghai, China).

### Yeast two hybrid assay and co-immunoprecipitation assay

The yeast two hybrid assay was performed as indicated in MATCHMAKER GAL4 two-hybrid system 3 (Clontech). The protein coding regions of genes used in this study were amplified from Guy11 cDNA with primer pairs listed in Table 1.1. Subsequent cloning of MoPtc1, Pmk1 and MoOsm1 was done in pGBKT7 bait vector and MoNbp1 in pGADT7 prey vector. These pairs were simultaneously co-transformed into *S. cerevisiae* strain AH109 as described by(62) together with a carrier DNA. The pGBKT7-53/pGADT7-T and pGBKT7-Lam/pGADT7-T vectors were used as positive and negative controls respectively. The emerged yeast colonies in Leu+ and Trp+ medium were isolated and cultured on SD-Trp-Leu-His-Ade selective medium containing specified concentration of X-α-gal.

For co-immunoprecipitation assay GFP-fusion-proteins from cellular extracts, were isolated and incubated with 20–30 μL of GFP-Trap agarose beads (ChromoTek, Germany) according to the manufacturer’s instructions. Proteins eluted from GFP-Trap agarose beads were analyzed by immunoblot detection with the anti-flag (Abmart), and anti-GFP (Abmart) antibodies respectively.

### Statistical analysis

The statistical analysis was performed using graph pad prism 7 and the data from the independent biological replicates was analyzed using a one-way ANOVA (and non-parametric) and the Dunnet multiple comparison test with 0.05 (95% confidence interval).

## Acknowledgements

This work was supported by National Natural Science Foundation of China (No.31301630 & No. 31601599) to Dongmei Zhang and Wenyi Yu. I would like to acknowledge Fujian government scholarship for financial aid as well as Dr. Guodong Lu, Dr. Jin-Rong Xu, Dr. Jie Zhou, Dr. Wenhui Zheng and Ms. Honghong Zhang for their resourceful ideas while running this project.

