## Supplemental Table1 & Figure for "The protein phosphatases MoPtc1 and MoPtc2 are induced during pathogen-host interactions and play synergistic roles in regulating MAPK pathways in *Magnaporthe oryzae*"

| **Name** | **Sequence 5' to 3'** |
| --- | --- |
| MGG_05207 AF | GCCAAAATGAGATACCAGAC |
| MGG_05207 AR | AGGGAACAAAAGCTGGGTACCCAGAGACGGTTGCAGAGACAC |
| MGG_05207 BF | GAATAGAGTAGATGCCGACCGCGGGTTGAGAAACAGCCCGCATAG |
| MGG_05207 BR | TTCAACGACCACGAAAGC |
| MGG_05207 OF | AAAACCACAGCCACTCCG |
| MGG_05207 OR | CGCTTGCTTGTCAAATCG |
| MGG_05207 UA | CGGTCGGTGGCGGTAGTGAT |
| MGG_01351 AF | TTGTGATTCTGTCGGTTC |
| MGG_01351 AR | TTGACCTCCACTAGCTCCAGCCAAGCCTTACGGTTGACTCCTGAG |
| MGG_01351 BF | GAATAGAGTAGATGCCGACCGCGGGTTTCCCCTACACCTTTGACCT |
| MGG_01351 BR | GCAATCTGAATCTCGTCCC |
| MGG_01351 OF | GGTGATGATGATGAGTTCT |
| MGG_01351 OR | ATCTTTGGTCCCTTTGTC |
| MGG_01351 UA | CATCTTTCCGAGGTGGCG |
| MGG_05207 Com-F | GAACAAAAGCTGGGTGAGAGGAGGCGCGTTTT |
| MGG_05207 Com-R | CTGCAGGCATGCAAGTTGAAGATGTGGCCGGTT |
| MGG_01351 Com-F | GAACAAAAGCTGGGTGTCGCAATACTCGGTCTT |
| MGG_01351 Com-R | CTGCAGGCATGCAAGGACCTTGATATCCTCGT |
| MoPTC1 AD-F | GTACCAGATTACGCTCATATGATGTTTGGCGGCTCCTC |
| MoPTC1 AD-R | ATGCCCACCCGGGTGGAATTCTTATGAAGATGTGGCCGGTT |
| MoPTC1 BD-F | TCAGAGGAGGACCTGCATATGATGTTTGGCGGCTCCTC |
| MoPTC1 BD-R | TCGACGGATCCCCGGGAATTCTTATGAAGATGTGGCCGGTT |
| MoNBP1 AD-F | GTACCAGATTACGCTCATATGATGTCTCGCGCCAATCC |
| MoNBP1 AD-R | ATGCCCACCCGGGTGGAATTCTTACCGCATAATTTCCTGG |
| MoPMK1 BD-F | TCAGAGGAGGACCTGCATATGATGTCTCGCGCCAATCC |
| MoPMK1 BD-R | TCGACGGATCCCCGGGAATTCTTACCGCATAATTTCCTGG |
| MoOSM1 AD-F | GTACCAGATTACGCTCATATGATGGCGGAATTCGTGCG |
| MoOSM1 AD-R | ATGCCCACCCGGGTGGAATTCTTATTGGCCGGTAAACT |
| MoOSM1 BD-F | TCAGAGGAGGACCTGCATATGATGGCGGAATTCGTGCG |
| MoOSM1 BD-R | TCGACGGATCCCCGGGAATTCTTATTGGCCGGTAAACT |
| H853 | GACAGACGTCGCGGTGAGTT |
| HG-F | GAATAGAGTAGATGCCGACCGCGGGTT |
| HG-R | TTGACCTCCACTAGCTCCAGCCAAGCC |
| MGG_05207 AF | GCCAAAATGAGATACCAGAC |
| MGG_05207 AR | GGAAATTGTAAGCGTTAATCTAGAGCGCGGTTGCAGAGACAC |
| MGG_05207 BF | GCATTCTGGGTAAACGACTCATAGGAGGAGAAACAGCCCGCATAG |
| MGG_05207 BR | TTCAACGACCACGAAAGC |
| MGG_01351 AF | TTGTGATTCTGTCGGTTC |

| **Name** | **Sequence 5' to 3'** |
| --- | --- |
| MGG_01351 AR | GGAAATTGTAAGCGTTAATCTAGAGCGTTACGGTTGACTCCTGAG |
| MGG_01351 BF | GCATTCTGGGTAAACGACTCATAGGAGTCCCCTACACCTTTGACCT |
| MGG_01351 BR | GCAATCTGAATCTCGTCCC |
| NEO/F | CGCTCTAGATTAACGCTTAC |
| NE/R | CCTGATGTTCTTCGTCCA |
| EO/F | GACAATCGGCTGCTCTGA |
| NEO/R | CTCCTATGAGTCGTTTACCCA |
| EO/F | GACAATCGGCTGCTCTGA |
| NR | CAATAGCAGCCAGTCCCT |

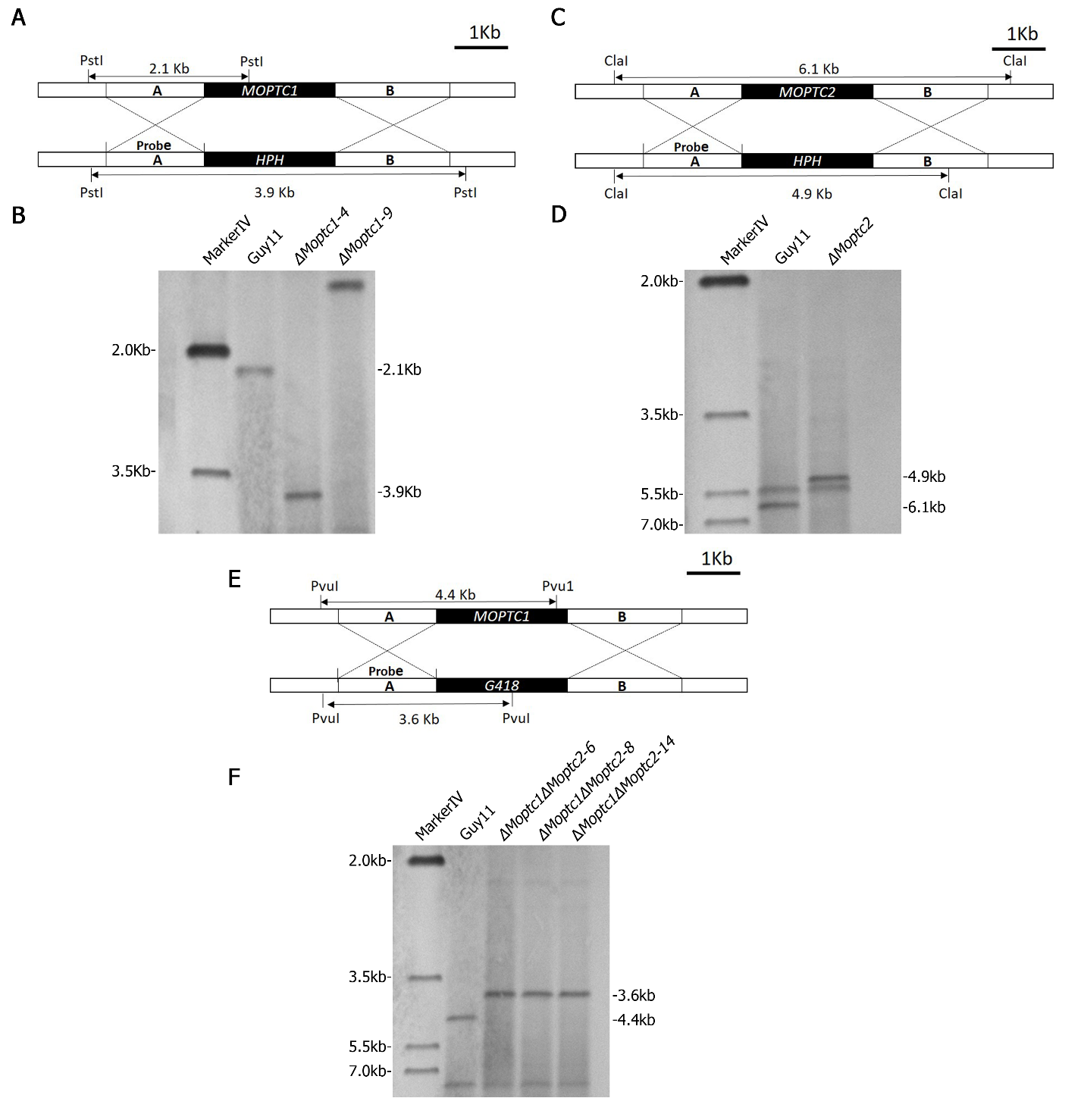

**Figure S1**. Targeted gene deletion for *∆Moptc1*, *∆Moptc3* and *∆Moptc1∆Moptc3* in *M. oryzae*.

**(A)**, **(C)** and **(E)** Schematic diagrams for MoPtc1, MoPtc3 and MoPtc1MoPtc3 disruptions via homologous recombination strategy. **(B)** and **(D)**  Southern blots confirmation results for *∆Moptc1*, and *∆Moptc3* replacements by single insertion of Hygromycin Phosphotransferase gene (Hph) in the respective ORF regions. **(F)** Southern blots confirmation results for *∆Moptc1∆Moptc3* replacements by single insertion of G418 resistance marker in Moptc1 mutant
